# Assessing Random Forest self-reproducibility for optimal short biomarker signature discovery

**DOI:** 10.1101/2023.03.29.534695

**Authors:** Christophe Poulet, Ahmed Debit, Claire Josse, Guy Jerusalem, Chloe-Agathe Azencott, Vincent Bours, Kristel Van Steen

## Abstract

Biomarker signature discovery remains the main path to develop clinical diagnostic tools when the biological knowledge on a pathology is weak. Shortest signatures are often preferred to reduce the cost of the diagnostic. The ability to find the best and shortest signature relies on the robustness of the models that can be built on such set of molecules. The classification algorithm that will be used is selected based on the average performance of its models, often expressed via the average AUC. However, it is not garanteed that an algorithm with a large AUC distribution will keep a stable performance when facing data. Here, we propose two AUC-derived hyper-stability scores, the HRS and the HSS, as complementary metrics to the average AUC, that should bring confidence in the choice for the best classification algorithm. To emphasize the importance of these scores, we compared 15 different Random Forests implementation. Additionally, the modelization time of each implementation was computed to further help deciding the best strategy. Our findings show that the Random Forest implementation should be chosen according to the data at hand and the classification question being evaluated. No Random Forest implementation can be used universally for any classification and on any dataset. Each of them should be tested for both their average AUC performance and AUC-derived stability, prior to analysis.

**Author summary:** To better measure the performance of a Machine Learning (ML) implementation, we introduce a new metric, the AUC hyper-stability, to be used in parallel with the average AUC. This AUC hyper-stability is able to discriminate ML implementations that show the same AUC performance. This metric can therefore help researchers in choosing the best ML method to get stable short predictive biomarker signatures. More specifically, we advocate a tradeoff between the average AUC performance, the hyper-stability scores, and the modeling time.

## Introduction

In the field of cancer care, biomarker screening helps clinicians in their decision making. Biomarker Signature Discovery (BSD) aims to identify a set of hundreds of variables, out of thousands, that will capture the molecular differences between the categories of patients to study. Short BSD focuses on the most relevant variables to build the shortest predictive signature for comfortable use in daily clinical routine. While the clinicians expect a manageable test with few variables, they also expect it to be robust enough to reduce the prediction error.

Short combinations of biomarkers, also called short signatures, can be easily transferred to the daily clinical routine. These short signatures can be used at low costs to diagnose a cancer subtype, predict treatment responses, or monitor the patient during the treatment [1, 2]. However, with more than 10,000 clinical trials, based on biomarkers and cancer, currently ongoing [3], only a few studies may be successfully transferred to the clinics, and fewer may impact the diagnosis practices as only a few biomarkers are clinically relevant yet [4]. Indeed, within the past clinical trials on Breast Cancer, only the Oncotype DX, MammaPrint, EndoPredict, Breast Cancer Index (BCI), and Prosigna (PAM50) multianalyte tests have been successfully transferred with their associated model of prediction [5]. Despite these few commercial successes, many publications are directly related to biomarkers [4], and the design of short, robust, and universal signatures of biomarkers predictive of a clinical state remains challenging.

Combinatory strategies are increasingly used to determine multivariate signatures [1, 6]. Machine Learning (ML) have gained popularity in this sense (missing) to create associated models [6–8]. Researchers often compare ML strategies upstream to determine the “best” approach to use, (missing) usually relying on the highest average AUC performance. However, such average AUCs are highly variable. Indeed, based on Gonzalez-Bosquet *et al*. Fig 2 A, nine ML methods including RF displayed AUCs between 0.6 and 0.75 [9]. Due to this variation, the average AUC is not an optimal criterion for assessing the best ML methodology.

Random Forest (RF) is among the most popular machine learning methods in bioinformatics and related fields. RF is an ensemble of classification or regression trees that was introduced by Breiman [10]. It is extensively applied on gene expression data because it copes with large *p* small *n* problems, it exhibits a relatively good accuracy, is robust to noise, and requires little parameter tuning. Moreover, RF is easy to use and the interpretation of the resulting models is facilitated since it is all about a suite of “if..else”-like decision rules. Since the original RF algorithm proposed by Breiman [10], several variations to RFs have been made available via the R Project for Statistical Computing, including orthogonal and oblique methods.

The current study aims at assessing RF strategies based on both the average AUC and the stability of the resulting AUC. Therefore, the question asked is how AUC stability can help deciding the best predictive RF implementation? We put this question in the context of short BSD identification and evaluation. In the BSD field, several applications of RFs exist [6, 9]. In this work, we focus on assessing the most stable RF method, from 15 implementations in R (see Table 1). Our study is driven by tumor *vs*. healthy in paired samples from The Cancer Genome Atlas (TCGA) database (RNAseq of BRCA, LUSC, and THCA cancers).

**Table 1.** Summary of the implementations used in this study. The R package column denotes the package version used in comparisons.

| Implementation | Type | OOB | R package |
| --- | --- | --- | --- |
| RandomForest | Orthogonal | ✓ | randomForest v4.6-12 [11] |
| RandomForestSRC | Orthogonal | ✓ | randomForestSRC v2.6.1 [12] |
| Ranger | Orthogonal | ✓ | ranger v0.10.1 [13] |
| cForest | Orthogonal | ✓ | partykit v1.1.1 [14] |
| Rborist | Orthogonal | ✓ | Rborist v0.1-8 [15] |
| ExtraTrees | Orthogonal |  | extraTrees v1.0.5 [16] |
| RandomUniformForest | Orthogonal | ✓ | randomUniformForest v1.1.5 [17] |
| RRF | Orthogonal | ✓ | RRF v1.7 [18] |
| WSRF | Orthogonal | ✓ | wsrf v1.7.17 [19] |
| iForest | Orthogonal |  | iRF v2.0.0 [20] |
| CCF | Oblique |  | ccf v1.0.0 [21] |
| PPForest | Oblique | ✓ | PPforest v0.1.1 [22] |
| obliqueRF | Oblique | ✓ | obliqueRF v0.3 [23] |
| RotationForest | Oblique |  | rotationForest v0.1.3 [24] |
| Rerf | Oblique | ✓ | rerf v1.0 [25] |

The conclusions are twofold. First, AUC-derived stability reveals the dataset dependency of an RF implementation. Second, based on two distinct scores, hyper-stability can highlight whether an RF implementation is signature or resampling dependent. Consequently, AUC stability provides a confidence score on top of the commonly used average AUC for the selection of the best RF implementation.

Additionally, the modelization time can further help discriminating between RF implementations with equal stability performance.

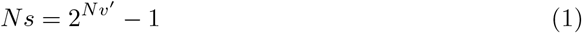

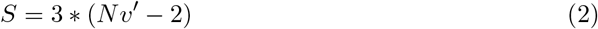

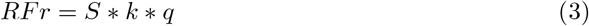

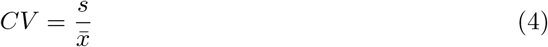

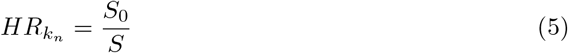

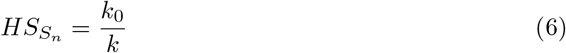

## Materials and methods

The description of the symbols used in the formulas is given in S1 Table.

### TCGA sample collection, normalization, and filtering

The TCGA database was screened to maximize the number of paired tumor-healthy samples in the cancer cohorts. TCGA clinical data were filtered to select cancers with the most abundant similar Histological subtype and patients with paired tumor-healthy samples. Subsequently, three TCGA datasets were targeted in the current study, the BRCA, the LUSC, and the THCA cancers. These three datasets were downloaded using the *TCGA2STAT* R-package [26]. Paired tumor-healthy samples were collected with RPKM normalization using the *tumorNormalMatch* function from the same R-package. The total number of primary variables was reduced based on their variance to *Nv* variables before the feature selection step, using the *Log Intensity variation* function of the BRB ArrayTools software (version 3.8.1) with the p-value parameter set to 0.001.

These filters were listed in S2 Table for the three datasets.

### Workflow of the comparative study

A graphical summary of our study comparing multiple RF implementations via hyper-stability assessment is given in Fig 1. Next, we explain each step in greater detail. In the following sections, we use the term “resampling rate” to refer to the percentage of data that goes to the training partition after a balanced random sampling from the original data.

**Fig 1.**
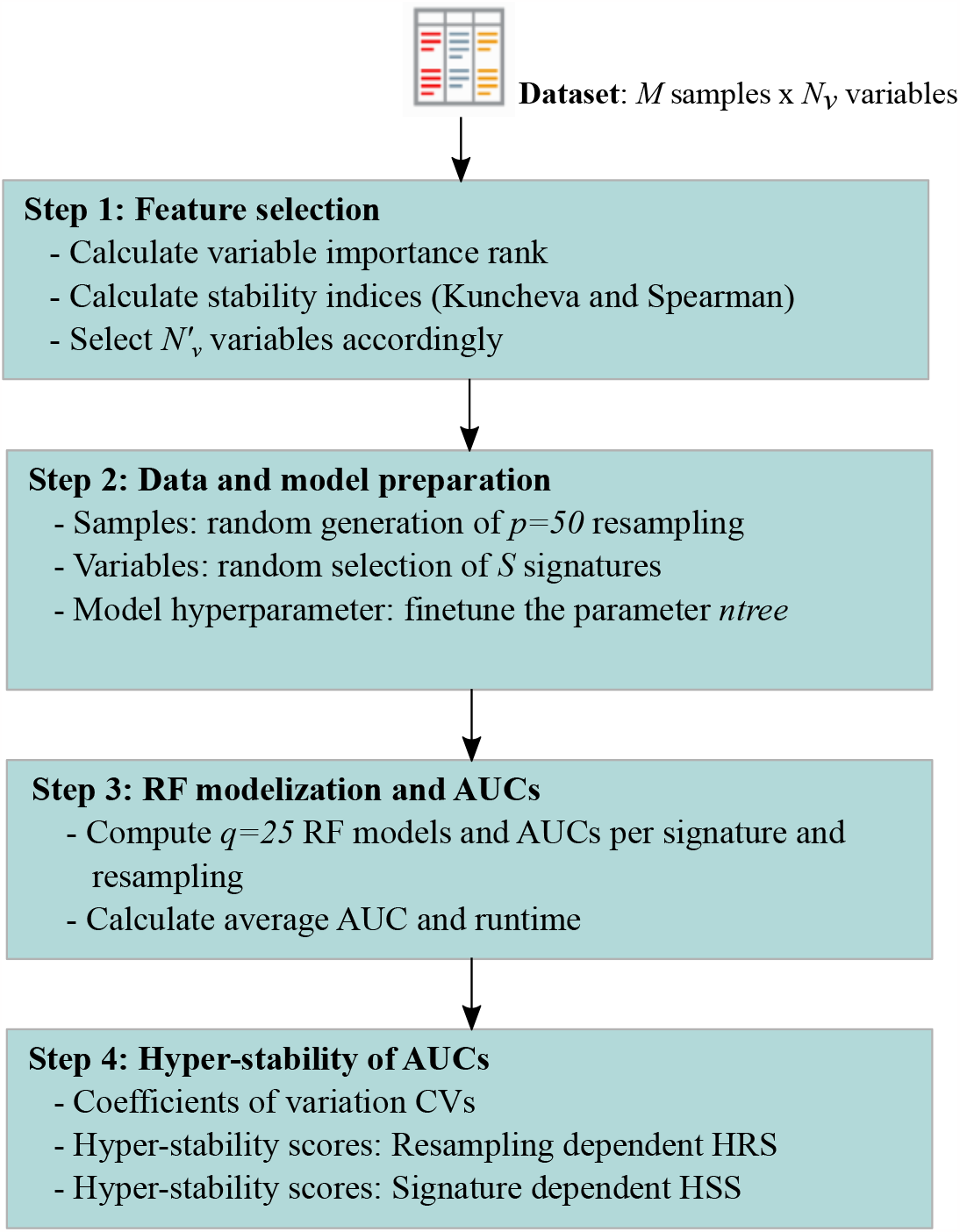
Comparative study flowchart: from feature selection to AUC hyper-stability. Overall procedure of AUC-based hyper-stability comparison. **Step 1-** Feature selection based on the variable importance rank and stability indices for the selection of the most important and most stable variables. **Step 2-** Preparation step including the generation of random partitions to be used as training and validation sets, the random selection of variable combinations, and the finetuning of the RF parameter *ntree*; **Step 3-** Modelization and validation: training and validation of 25 RF models per signature and resampling partition, with recorded runtime; **Step 4-** AUC-based hyper-stability scores HRS and HSS to complement average AUCs across signatures and samples

### Step 1: Feature selection

Feature Selection (FS) followed rank-based overlap and correlation principles inspired by Alelyani et al [27]. FS was used to determine the minimum number of variables *Nv*^*′*^ to be kept for downstream analysis. Rank-based variable importance was used and was calculated based on a combination of the mean decrease in GINI (MDG), and mean decrease in Accuracy (MDA) of the RF algorithm. The threshold used for the selection of *Nv*^*′*^ was determined using stability indices Kuncheva and Spearman. The entire methodology is provided in supplementary (S1 File).

### Step 2: Data and model preparation

#### Fine tune the parameter *ntree*

For all RF methods that implemented the out-of-bag principle (see Table 1) the out-of-bag error (OOBerr) was computed based on *Nv*^*′*^ variables and increased values of *ntree* ∈ {10, 20, .., 1200}. Similarly, a total of *k* = 50 balanced random case/control partitions were generated using a resampling rate of *p* = 0.9, and *q* = 25 intrinsic RF models per partition. Subsequently, 50 × 25 = 1,250 models and 1,250 OOBerr values were obtained for each value of *ntree*. The OOBerr was averaged over the *ntree* value and plotted as a function of *ntree* for each RF method. The minimal number of trees *Nt* = 500 was obtained when the OOBerrs were optimized and stabilized for the tested implementations. This value of *Nt* is sufficiently acceptable from all the RF implementations including those not implementing the out-of-bag principle.

#### Obtain potential signatures of max length *Nv*^*′*^

The total number of possible signatures *Ns* was defined by Eq 1. These signatures contained a different number of variables, from size 1 to size *Nv*^*′*^. A random selection of three signatures from size 2 to size *Nv*^*′*^ *−* 1 was made (see S3 Table). A total of *S* signatures were therefore selected for the comparison using Eq 2.

#### Define learning and validation sets

A total of *k* = 50 random training partitions were generated from the original dataset, using a resampling rate *p* = 0.5 for downstream analysis. The function *createDataPartition* from the R-cran package *caret* [28] *was used to create these partitions*.

### Step 3: RF Models and AUCs

*Each RF implementation was computed with the ntree* parameter *Nt* for a total of *RFr* times based on Eq 3; where *S* is the number of signatures based on Eq 2, *k* is the number of random partitions, and *q* is the number of models generated per partition.

For the current article each RF implementation was sent to an individual computational node for learning and validation, with *k* = 50 and *q* = 25. Each node was, therefore, handling *RFr* computations, which resulted in *RFr* models and *RFr* AUCs. Metrics for each model validation were also computed with the R-package *MLmetrics* [29]. *The time was measured before and after each modelization*.

*To get the average AUC of an implementation, the AUC was averaged across RFr* runs. Similarly, the time to process the model was averaged across *RFr* runs to get the average runtime of the implementation.

To measure the runtime, we selected computational nodes having the same characteristics.We used the nodes from the CECI’s Dragon1 cluster hosted by the UMons university Belgium, which provided 416 CPU distributed on 26 nodes and 128*GB* of RAM. The CPUs used were *SandyBridge* processors of 2.60*GHz*.

### Step 4: Hyper-stability

For each RF implementation’s resampling-signature combination, the coefficient of variation (CV) of the *q* AUCs was computed. In the current article, *q* = 25. This CV measures the average variability of AUCs around the mean AUC, defined by Eq 4, where *s* is the sample standard deviation and 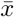 is the sample mean. It lies between 0 and 1.

The hyper-stability scores were computed based on *CV* == 0 values. Two novel hyper-stability scores were created, a resampling dependent (*HRS*) and a signature dependent (*HSS*) score. For *HRS*, signatures with *CV* == 0 (in total *S*_0_) were averaged across the total number of signatures *S* tested for a resampling *k*_*n*_ and was named *HR*, as displayed by Eq 5. The mean of non-zero *HRs* across all the resamplings was then used to create the *HRS* score, which describes the method on a resampling base. The *HSS* averages the number of resampling with *CV* == 0 denoted *k*_0_ across the total number of resampling *k* tested for a signature and was named *HS*, as displayed by the Eq 6. The mean of non-zero *HSs* was then used to create the *HSS* score, which describes the method on a signature base.

We used R version 3.3.1 for all the analyses detailed in the current study. Table 1 listed all RF implementation R-packages and their versions used in the current study.

## Results

### Hyper-reproducibility of the AUCs

To assess each RF implementation’s modelization, we identified which signature-resampling combination could produce the same AUC, using the Coefficient of Variation (CV). A dot-matrix Fig 2 displayes all the LUSC dataset *CV* == 0, obtained for all combined signature-resampling of each RF implementation. Then we classified the RF implementations into three groups based on lack of reproducibility. i) The signature dependent group (type A) contained only PPForest, which displayed multiple blank rows; ii) The resampling dependent group (type B) encompassed ccf, rerf, randomForest, iForest, and ranger, which presented numerous empty columns; iii) The signature-resampling group (type C) was composed of the remaining implementations where no trend neither in signature nor in resampling could be found. Aside from our classification, RRF showed only a few stable signature-resampling combinations and was therefore unclassified. RF implementations switched between groups depending on the dataset under study (results not shown).

**Fig 2.**
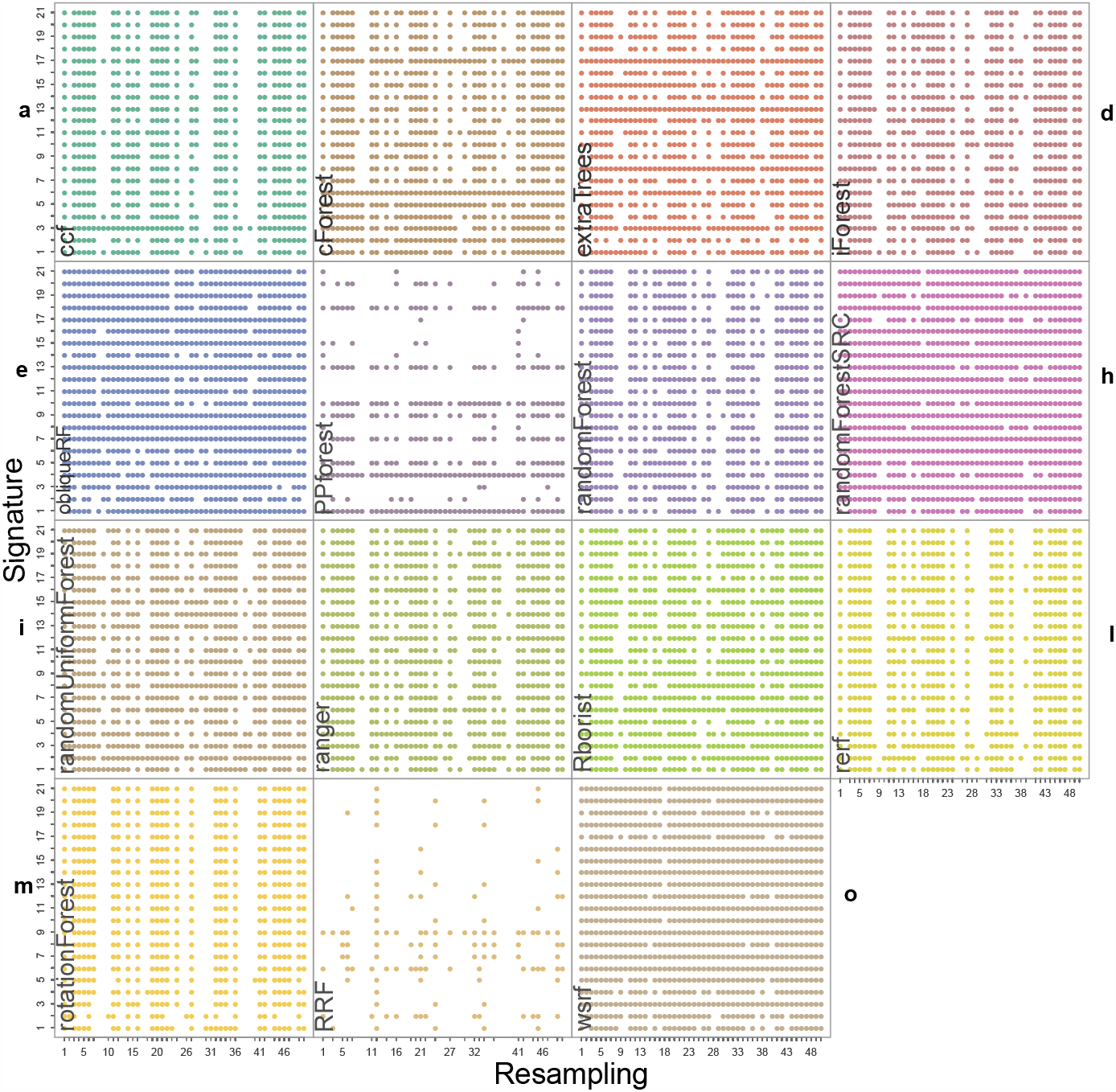
Dot-matrix of AUCs coefficient of variation equal to 0 for all RF implementations. Coefficient of variation equal to 0 for *q* = 25 AUCs obtained for each signature-resampling combination per RF implementation for the LUSC dataset. Each dot corresponds to 25 equal AUCs (*CV* == 0) for a signature-resampling combination. No dot correponds to a variation in the 25 AUCs (*CV >* 0) for a signature-resampling combination.

### Hyper-stability score

We computed the hyper-stability scores for resampling (HRS) and signature (HSS) to further compare the RF implementations. Dot-marix Fig 2 was used to calculate HRS and HSS (see Fig 3.a for an example how to calculate HRS and HSS from the dot-matrix). A bar chart to display these HRS (green bars) and HSS (purple bars) for the LUSC is given in (Fig 3.b). The type A group PPforest method shows a combined score below 0.4. The type B implementations shows almost a similar values of HRS and HSS with a combined value *≥*0.65. Furthermore, among the type C group, only obliqueRF, randomForestSRC, and wsrf obtained a combined score around 0.9, while the remaining implementations obtained a score below 0.8. We observed similar trends for the BRCA dataset. However, the THCA dataset displayed HRS and HSS scores below 0.4 for all implementations. More specifically, for the LUSC dataset, **good** scores (]0.8, 1]) were obtained for obliqueRF, randomForestSRC and wsrf; **moderate** scores ([0.4, 0.8]) for rotationForest, ccf, cForest, extraTrees, iForest, randomForest, RUF, Rborist, ranger, Rborist, rerf, and rotationForest; and **poor** scores ([0, 0.4[), for RRF and PPforest. Interestingly, the poorly performing group implementations persisted across datasets (LUSC, BRCA, and THCA). The good and moderate RF implementations were inconsistent across datasets. These results underlined the dataset dependency of the RF implementations studied here.

**Fig 3.**
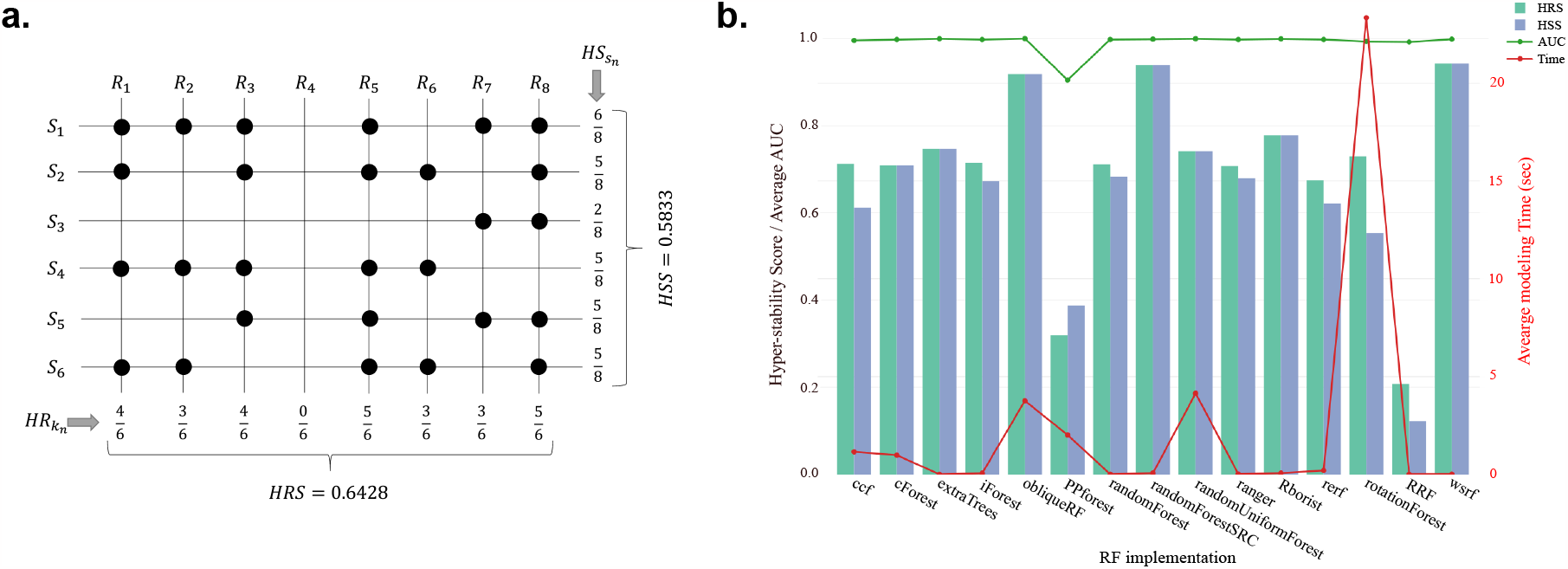
Random Forest hyper-stability scores. a) Example of calculating the HR (Eq 5), the HS (Eq 6), HRS, and HSS. b) HRS score (green) and HSS score (purple) are displayed as barplots for each RF implementation, using the left scale of the graph. Average AUC of each RF implementation is reported with a green line using the left scale of the graph. Average time to process the modelization is reported with a red line using the right scale of the graph.

### Average AUC and time of modelization

We calculated the average AUC obtained from *RFr* modelizations. We also calculated the average time to process a model. Fig 3.b displays this average AUC (green line) and time (red line) obtained by each RF implementation for the LUSC dataset. Except for PPforest and RRF, the average AUC was equal to 1 for all the RF implementations. With an average time of 23.3 sec to process a model, rotationForest was the slowest method. RF implementations ccf, cForest, obliqueRF, PPForest, and randomUniformForest processed the models with an average time between 1 and 4.1 seconds. The remaining RF implementations processed the models with an average time below 0.2 seconds. Similar trends were observed for the BRCA and THCA datasets but with a higher modelization time (data not shown).

### Identification of probable causes of hyper-stability scores impairment

#### Algorithm taxononomic classification

We further assessed whether the hyper-stability scores varied according to the RF implementation taxonomy previously described by Pretorius *et al*. [30]. Based on the 15 RF implementations, the following four criteria could be derived from this taxonomy: number of layers of randomization modification; transformation or projection of the dataset; non-exhaustive search, and deterministic modifications.

Taking the randomForest original implementation as a reference, we assessed these four criteria’ impact on the HRS and HSS scores. Adding or removing layers of randomization did not improve the scores. Similarly, non-exhaustive search methods did not drastically change the scores. Indeed, extraTrees or RUF did not show the highest HRS and HSS scores. Besides, data transformation or projection might have an impact on the scores. We observed a decrease of the HRS and the HSS scores for rotationForest, PPforest and ccf, as well as an increase of these scores for rerf for the BRCA dataset or obliqueRF for the LUSC dataset. The deterministic modification might also impact the scores, especially for RRF, PPforest, cForest, rerf, and wsrf. Nevertheless, no impact of this deterministic modification could be detected with the scores obtained by RUF, ramdomForest, SRC, Rborist, and obliqueRF. Importantly, no common impact could be linked to Pretorius et al. taxonomy to the HRS and HSS scores.

#### Dataset properties

To assess whether the dataset’s size may impair the hyper-stability scores, we further explored the variable-sample ratio (*number of variables / number of samples*) in the datasets. With a data perturbation of 0.5, 0.5 *\** 96 = 48 samples were randomly selected in each LUSC resampling, leading to a variable-sample ratio of 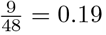. By decreasing the data resampling rate to 0.9, the variable-sample ratio decrease to 0.1, which resulted in higher HRS and HSS scores for all implementations (data not shown). Despite the same number of samples in the THCA dataset, this ratio reached 0.77 for a resampling rate of 0.5, leading to HRS and HSS scores around 0.4 for all the implementations. Again, the scores could be increased over 0.8 by switching to a variable-sample ratio of 0.44 at a resampling rate of 0.9. Besides, with 28 variables and 182 samples, the BRCA variable-sample ratio reached 0.3 for a resampling rate of 0.5. Consequently, HRS and HSS scores were likely to depend on the variable-sample ratio, where good scores (*>* 0.8) were obtained with a low variable-sample ratio (*<* 0.5).

To evaluate whether the resampling could impair an RF implementation’s hyper-stability, we quantified the resampling with few *CV* == 0. Using the dot matrix based on LUSC, Fig 2, we observed that resampling 37 and 38 were struggling to stabilize the AUC for most RF implementations. Such a lack of stability on the LUSC dataset was particularly significant for the ccf, PPforest, rerf, rotationForest, and RRF implementations. We made similar observations for BRCA’s resampling 6 and 34 and THCA’s resampling 6 to 10. Consequently, the presence of these problematic resamplings might contribute to the decrease of the hyper-stability scores.

To better understand the HRS and HSS variabilities between the datasets, we looked into gene connectivity. Differences in connectivity might explain the modelization performances of machine learning. Using WGCNA [32] on the 7031 genes in common between the tumor samples of the three filtered datasets, we seek for modules with highly correlated genes conserved between the three datasets. Thus, we compared *BRCA >> LUSC, BRCA >> THCA*, and *LUSC >> THCA* (S1 Fig) *We found that 15 modules among 25 showed low preservation for BRCA >> LUSC*. When performing a module preservation analysis, genes with a similar function tend to cluster together according to a phenotype or a disease [31]. These modules were located between no preservation (blue line, *Zsummary* = 2) and very weak preservation (green line *Zsummary* = 10) region. These modules also had the lowest median rank statistic, meaning their observed preservation statistics tend to be the lowest among the other modules. Moreover, 14 highly connected genes within the grey60 module of BRCA lost their connectivity within the LUSC network (S2 Fig). Similarly, for the *BRCA >> THCA* case, we observed that 15 modules among 25 were lowly preserved in the THCA samples, and 20 modules out of 24 showed weak preservation for the *LUSC >> THCA* case. These results underlined that the functional connectivity was not conserved between the three datasets, and further contribute to explain differences in HRS/HSS scores.

## Discussion

In this study, we tested the stabilities of 15 different RF modelizations over three distinct cancer datasets. These datasets were composed of paired tumor-healthy samples. We compared RF modelizations for AUC hyper-stability scores and runtime, and investigated drivers of (in)stability.

We assessed RF’s hyper-stability based AUC-derived HRS and HSS scores for 15 RF implementations. The AUC has become a common measure to determine the accuracy of classification models. Nevertheless, it only evaluates how much the classification model can discriminate the classes. The AUC could, therefore, be misleading and might suffer from the following drawbacks [33]: AUC i) ignores the probability values of the samples; ii) includes less interesting regions on the ROC plot; iii) does not reflect the intended use of the model; iv) does not provide information about the spatial distribution of model errors. Besides, the AUC might be insensitive to strongly-associated disease features added to the model [34]. Moreover, AUC displays a high dispersion, especially for imbalanced or small sample sets [35]. AUC alternatives can provide useful measures of performance for prognostic models [35]. Examples are the Pietra index and the standardized Brier or scaled Brier scores. These alternatives should be considered for future calculation of the hyper-stability scores.

In this work, we built on AUC for different scenarios of resampling and signature combinations to derive hyper-stability scores HRS and HSS. The proposed methodolgy tried to assess RF inherent randomness while keeping the external randomness under control. While we measured the exact same AUC from 25 models, our system was not deterministic, as defined by Padhye *et al*. [36]. Indeed, rule extraction showed that the genes, the thresholds, and the number of steps used could differ in each model (see supporting information section (S1 File)). Consequently, our system kept the intrinsic randomness of the RF implementations and could not be considered deterministic. Good HRS and HSS scores were obtained in our study, except for the THCA dataset at a perturabtion rate of 0.5. With a perfect balance between tumor and healthy samples, the small sample size might explain THCA’s low performances. Nevertheless, the LUSC dataset displayed good hyper-stability scores with the same number of samples.

Specific characteristics of the RF algorithm and nature of the application data are the main drivers to model performance (S4 Fig), which we discuss next.

- **Algorithm characteristics:** RF algorithms may differ from Breiman’s original implementation in their randomization and deterministic components. Except for the tree selection and the ensemble compilation, the 15 chosen RF implementations covered all the taxonomy criteria listed by [37]. The following characteristics could therefore impact AUC performances:
  i. The number of trees; A model with more trees is better [38]. Our results were based on 500 trees for each RF implementation. Indeed, except for RRF, the resulting OOBerr stabilized for most RF implementations after 500 trees (S3 Fig c). RRF struggled to stabilize the OOBerr after 500 trees, which might explain its poor hyper-stability over all the datasets.
  ii. The sources of randomization encompass; selecting samples, sampling the features, and selecting the splitting point [37]. The randomization component deals with the insertion or deletion of randomization layers and the modification of the random sampling procedure.
  iii. The deterministic modifications, which encompass; oblique or orthogonal splits, impurity measure, and penalization [37]. The deterministic component deals with the tree construction, data transformation, type or rules to split, impurity measure, or variable penalities. No relationships were found between the hyper-stability scores, the sources of randomization, and the deterministic modifications. However, “good”, “moderate”, and “poor” groups could be derived from the hyper-stability scores. Interestingly, only the “poor” group contained dataset-independent RF implementations, RRF and PPForest. For the purpose of the current study, we used an RF-based FS coupled with the Kuncheva and Spearman indexes to check the overlap and the correlation of the variable ranking. The aim was to get the minimal set of important and stable variables before performing the comparison. The FS did not favor any RF implementation method, it may negatively impact the “poor” group. For example, RRF uses regularized selected variables during FS. Further work is thus needed to study how such regularization may affect the FS used here. This work might be done for RRF and PPforest using the LDA-based projection pursuit index to identify projections that separate classes [39]. *Conversely, the Good* and the *Moderate* groups contained non-fixed RF implementations regarding the dataset studied. Indeed, each RF implementation resulted in different HRS and HSS scores when facing another dataset, and thus might be classified as *Good* or *Moderate*.
- **Dataset characteristics:** Although the RF versatility between *Good* and *Moderate* groups could not be ignored, it might be linked to the training set. Indeed, the selected samples and variables should provide enough information to the model to fully recognize the patterns. A low variable-sample ratio could be critical to produce models with a high average AUC [40] and a good hyper-stability. The following dataset characteristics could, therefore, impact the AUC.
  i. The balance between the categories: While the three datasets used in the current study were perfectly balanced between the tumor and the healthy classes, extremely imbalanced classes could harm RF behaviors [41]. To apply our methodology on non-balanced datasets, we recommend the use of balanced or weighted RF implementations. Chen *et al*. proposed balanced and weighted RF, where balanced RF will force to deal with equally sized classes, and the weighted RF were based on cost-sensitive learning. For such skewed datasets, the precision-recall curve (PR-curve) and the weighted-AUC should be preferred over the ROC-curve and the AUC [42, 43].
  ii. The size of the training set: The number and the heterogeneity of the samples could be an essential source of instability during the biomarker selection, leading the training set to be more or less attractive for the RF [44–46]. Subsequently, thousands of samples were recommended to reach good Kuncheva and Spearman scores [40]. However, our results achieved acceptable scores with the FS and only 192, 96, and 98 samples, meaning that a good apriori on the samples could circumvent this issue. Nevertheless, few resamplings displayed a low number of *CV* == 0 across RF implementations but impacted them similarly. A good apriori on the sample classes could, therefore, lead to heterogeneous random resamplings. Thus, such heterogeneity could impact the FS and the hyper-stability scores [27]. Interestingly, the variable-sample ratio appeared to be more related to the variability observed for the hyper-stability scores. By keeping the variable-sample ratio below 0.5, we obtained good hyper-stability scores (*>* 0.8) for the implementations. For example, we observed that the AUC reproducibility decreased along with the signature size for the THCA dataset, which was the only dataset that displayed a high variable-sample ratio and an average of low hyper-stability scores. Also, the length of a signature might impact the *mtry* parameter and might impair their hyper-stability [47]. However, by decreasing the data perturbation of the training step to 0.9, the RF implementations reached good hyper-stability scores while keeping the variable-sample ratio below 0.5. Therefore, reducing the *Nv*^*′*^ or the data perturbation for the THCA dataset could keep the variable-sample ratio below 0.5, leading to higher hyper-stability scores. Remarkably, we repeated all the modelizations on three different sets of increasing signature sizes for each dataset, and the same hyper-stability scores (avg *SD* = 0.01) were obtained each time for each RF implementation. Subsequently, the HSS and HRS scores were more tied to the models and their overfitting when the variable-sample ratio was kept under 0.5.
  iii. The feature connectivity: By grouping the genes into modules of highly co-expressed genes, we could assess the FS bias occurring when highly-connected genes are selected. Our results demonstrated the difference between the three datasets in the gene-gene connectivity across the tumor samples, meaning that the functional information was different between the datasets. While 60% of the modules displayed a low preservation from BRCA to LUSC (15/25) or from BRCA to THCA (15/25), 83% of the modules displayed a weak preservation from LUSC to THCA (20/24). Such a difference might explain the variation also observed in AUC-hyper-stability between the datasets. Nevertheless, more work is needed to find a causative link between such connectivity and either the AUC-hyper-stability or the data perturbation by performing module preservation between the training partitions.
  iv. The correlations within signatures: Datasets with many correlated variables may have created misleading feature rankings [48]. Very few strong correlations were found within BRCA signatures, while this dataset displayed excellent hyper-stability scores. Subsequently, the correlations between variables within the signatures could not explain lower hyper-stability scores observed for both LUSC and THCA datasets.
  v. The variability of the dataset: With a high entropy, datasets tend to be more sensitive to perturbation, which results in different AUC performance. This is often the case of small datasets like biological data. In this study, the FS allowed us to maximize the class separability and separate the samples according to the tumor or healthy groups. However, while a perfect separation was observed for the BRCA and LUSC datasets, it was close to perfect for the THCA dataset. Indeed, all but four THCA samples were linked to their respective class. These few crossing-class samples might explain both the high-number of THCA variables after the FS and the low AUC hyper-stabilities. Further work is therefore needed to assess if the class separability could impair the AUC hyper-stability.

Intuitively, as defined in the current study the HR score relies on the robustness of the signature tested because the HR score allowed us to measure the AUC-performance reproducibility of a signature over different resampling. On the other hand, the HS score can refer to the signature multiplicity invoked in [49], because the HS score allowed us to assess the performance of multiple alternative signatures on one resampling partition.

## Conclusion

In the current study, we demonstrated the importance of measuring RF implementations’ hyper-stability in the context of short BSD. We reinforced the message that no RF implementation should be used blindly for classification and on any datasets. Instead, each should be tested for their AUC performance and AUC-derived hyper-stability before the analysis. While the AUC-derived hyper-stability could reveal the dataset dependency of an RF implementation, it could also identify the origin of such reliance, telling whether an RF implementation is signature or resampling dependent. Therefore, the hyper-stability scores measured a trustable difference that should be taken into account while comparing the RF implementations. Moreover, the modelization time could further help discriminate RF implementations with equal hyper-stabilities. Consequently, the AUC hyper-stability and the modelization time reinforce the average AUC message and guide the researcher towards the best RF strategy for short biomarker signature discovery or other fields.

## Supporting information

**S1 Fig. Evaluation of module preservation between tumor samples of the three datasets:** a- *BRCA >> LUSC*; b- *BRCA >> THCA*; and c- *LUSC >> THCA*. Horizontal dashed lines on the preservation Zsummary plots delimit weak preservation zone. Modules located above the blue line are not preserved.

**S2 Fig. Example of the change in connectivity between genes within the grey60 (***BRCA >> LUSC* **analysis)** a- The 14 most connected genes within the grey60 module in BRCA tumor samples are selected, and b- the corresponding connections in the LUSC network are calculated and plotted. The network is generated using VisANT software.

**S3 Fig. Feature selection on the BRCA-TCGA dataset**. a) Stability indexes for the FS were calculated for an increasing cardinality from 10 to 200 with a step of 10. The minimal number of stable variables (30 for the BRCA-TCGA dataset) was set as the first local maxima observed on the Kuncheva index (vertical blue line); b) The rank distribution of the top 200 variables. For each variable, all the 1250 ranks were displayed as a boxplot. The variables were then ordered based on their average rank. The minimal number of important variable obtained from figure a was reported as a vertical blue line. The adjustement to this number was reported here as a vertical red line; c) The prediction error (OOB error) was calculated for the methods implementing the OOB concept. The parameters used to compute this OOB errors were based on results obtained with figure a, b. The error was stable after 500 trees (vertical blue line) for all the implementations.

**S4 Fig. AUC performance impacting factors** Diagram showing factors impacting the AUC performance of an RF algorithm. Some factors concern dataset characteristics, others relate on the construction of RF algorithm. Both randomization sources and deterministic modifications components are used to construct RF variants (inspired from [37])

**S5 Fig. PCA of samples before and after the Feature Selection** PCA projection of the 2 first principal components for a) BRCA; b) LUSC, and c) THCA datasets. For each graph, the tumor samples are in red and their paired healthy tissues are in green. The Feature Selection increases the separatibility of the samples according to their respective class (tumor or healthy).

**S1 Table. Description of different symbols used in the formulas**.

**S2 Table. Summary of the results of step 1 and step 2 on TCGA datasets**. The row ‘Selected Signatures’ shows the number of combinations retained for the comparison. The statistics value is the combined value obtained from the Kunchva and the Spearman statistics. The dispersion is the optimal value obtained from the boxplots of the variable ranks.

**S3 Table. Table summarizing the signatures selected per dataset:** a) BRCA, b) LUSC, c) THCA

**S4 Table. Rules randomly extracted from two randomForest models trained on the same partition and the same signature for the LUSC dataset**.

**S5 Table. Three sets of rules randomly extracted from two randomForest models**. The RF models were trained on the same partition and the same signature for the BRCA, the LUSC, and the THCA datasets.

**S6 Table. Supplementary Range of CVs and number of** *CV* == 0 **observed for LUSC dataset**.

**S1 File. Supplementary Materials** Supplementary methods, results, and discussion

## Acknowledgements

We are extremely grateful for the logistic and organisational support that this study has received. Computational resources have been provided by the Consortium des Équipements de Calcul Intensif (CÉCI).

## Notes

### Competing Interest Statement

The authors have declared no competing interest.

## References

1. Enroth S, Berggrund M, Lycke M, Broberg J, Lundberg M, Assarsson E, et al. High throughput proteomics identifies a high-accuracy 11 plasma protein biomarker signature for ovarian cancer. Communications Biology. 2019;2(1):1–12. doi:10.1038/s42003-019-0464-9.

2. Tarhini A, Kudchadkar RR. Predictive and on-treatment monitoring biomarkers in advanced melanoma: Moving toward personalized medicine. Cancer Treatment Reviews. 2018;71(March):8–18. doi:10.1016/j.ctrv.2018.09.005.

3. NIH. Clinical Trials on Cancer and Biomarkers; 2019. Available from: https://clinicaltrials.gov/ct2/.

4. Selleck MJ, Senthil M, Wall NR. Making Meaningful Clinical Use of Biomarkers. Biomarker Insights. 2017;12:1–7. doi:10.1177/1177271917715236.

5. Duffy MJ, Harbeck N, Nap M, Molina R, Nicolini A. ScienceDirect Clinical use of biomarkers in breast cancer : Updated guidelines from the European Group on Tumor Markers. European Journal of Cancer. 2017;75:284–298. doi:10.1016/j.ejca.2017.01.017.

6. Frères P, Wenric S, Boukerroucha M, Fasquelle C, Thiry J, Bovy N, et al. Circulating microRNA-based screening tool for breast cancer. Oncotarget. 2015;7(5):5416–5428. doi:10.18632/oncotarget.6786.

7. Marcišauskas S, Ulfenborg B, Kristjansdottir B, Waldemarson S, Sundfeldt K. Univariate and classification analysis reveals potential diagnostic biomarkers for early stage ovarian cancer Type 1 and Type 2. Journal of Proteomics. 2019;196(March 2018):57–68. doi:10.1016/j.jprot.2019.01.017.

8. Boylan KLM, Geschwind K, Koopmeiners JS, Geller MA, Starr TK, Skubitz APN. A multiplex platform for the identification of ovarian cancer biomarkers. Clinical Proteomics. 2017;14(1):1–21. doi:10.1186/s12014-017-9169-6.

9. Bosquet JG, Newtson AM, Chung RK, Thiel KW, Ginader T, Goodheart MJ, et al. Prediction of chemo-response in serous ovarian cancer. Molecular Cancer. 2016;doi:10.1186/s12943-016-0548-9.

10. Breiman L. Random Forests. Machine Learning. 2001;45:5–32.

11. Liaw A, Wiener M. Classification and Regression by randomForest. R News. 2002;2(3):18–22.

12. H I, B KU. Random survival forests for R. R News. 2007;7(2):25–31.

13. Wright MN, Ziegler A. ranger: A Fast Implementation of Random Forests for High Dimensional Data in C++ and R. Journal of Statistical Software. 2017;77(1):1–17. doi:10.18637/jss.v077.i01.

14. Torsten Hothorn AZ. partykit: A Modular Toolkit for Recursive Partytioning in R. Journal of Machine Learning Research. 2015;16:3905–3909.

15. Seligman M. Rborist: Extensible, Parallelizable Implementation of the Random Forest Algorithm; 2019. Available from: https://CRAN.R-project.org/package=Rborist.

16. Simm J, de Abril IM, Sugiyama M. Tree-Based Ensemble Multi-Task Learning Method for Classification and Regression; 2014. Available from: http://CRAN.R-project.org/package=extraTrees.

17. Ciss S. randomUniformForest: random Uniform Forests for Classification, Regression and Unsupervised Learning; 2015. Available from: http://CRAN.R-project.org/package=randomUniformForest.

18. Deng H, Runger G. Feature selection via regularized trees. Proceedings of the International Joint Conference on Neural Networks. 2012;doi:10.1109/IJCNN.2012.6252640.

19. Zhao H, Williams GJ, Huang JZ. wsrf: An R Package for Classification with Scalable Weighted Subspace Random Forests. Journal of Statistical Software. 2017;77(3):1–30. doi:10.18637/jss.v077.i03.

20. Basu S, Kumbier K. iRF: iterative Random Forests; 2017. Available from: https://CRAN.R-project.org/package=iRF.

21. Dobler J, Feuerriegel S. ccf: Canonical correlation forest; 2016. Available from: https://github.com/jandob/ccf.

22. da Silva N, Lee EK, Cook D. PPforest: Projection Pursuit Classification Forest; 2018. Available from: https://CRAN.R-project.org/package=PPforest.

23. Menze B, Splitthoff N. obliqueRF: Oblique Random Forests from Recursive Linear Model Splits; 2012. Available from: https://CRAN.R-project.org/package=obliqueRF.

24. Ballings M, Van den Poel D. rotationForest: Fit and Deploy Rotation Forest Models; 2017. Available from: https://CRAN.R-project.org/package=rotationForest.

25. Browne J, Tomita T. rerf: Randomer Forest; 2019. Available from: https://CRAN.R-project.org/package=rerf.

26. Wan YW, Allen GI, Anderson ML, Liu Z. TCGA2STAT: Simple TCGA Data Access for Integrated Statistical Analysis in R; 2015. Available from: https://CRAN.R-project.org/package=TCGA2STAT.

27. Alelyani S, Liu H, Wang L. The Effect of the Characteristics of the Dataset on the Selection Stability. In: 2011 IEEE 23rd International Conference on Tools with Artificial Intelligence; 2011. p. 970–977.

28. Kuhn M. Building Predictive Models in R Using the caret Package. Journal of Statistical Software, Articles. 2008;28(5):1–26. doi:10.18637/jss.v028.i05.

29. Yan Y. MLmetrics: Machine Learning Evaluation Metrics; 2016. Available from: https://CRAN.R-project.org/package=MLmetrics.

30. Pretorius A, Bierman S, Steel SJ. A meta-analysis of research in random forests for classification. In: 2016 Pattern Recognition Association of South Africa and Robotics and Mechatronics International Conference (PRASA-RobMech). IEEE; 2016.

31. Langfelder P, Luo R, Oldham MC, Horvath S. Is my network module preserved and reproducible? PLoS Computational Biology. 2011;7(1). doi:10.1371/journal.pcbi.1001057.

32. Langfelder P, Horvath S. WGCNA: An R package for weighted correlation network analysis. BMC Bioinformatics. 2008;9. doi:10.1186/1471-2105-9-559.

33. Lobo JM, Jiménez-valverde A, Real R. AUC : a misleading measure of the performance of predictive distribution models. Global Ecology and Biogeography. 2008; p. 145–151. doi:10.1111/j.1466-8238.2007.00358.x.

34. Wu Yc, Lee Wc. Alternative Performance Measures for Prediction Models. PLoS ONE. 2014;9(3). doi:10.1371/journal.pone.0091249.

35. Hanczar B, Hua J, Sima C, Weinstein J, Dougherty ER. Small-sample precision of ROC-related estimates. Bioinformatics Advance. 2010; p. 1–9.

36. Padhye S, Sahu RA, Vishal S. Introduction to cryptography; 2018.

37. Pretorius A, Bierman S, Steel SJ. Advances in Random Forests with Application to Classification. Books. 2016;1(December):1.

38. Probst P, Boulesteix AL. To Tune or Not to Tune the Number of Trees in Random Forest. Journal of Machine Learning Research. 2018;18:1–18.

39. Lee YD, Cook D, Park JW, Lee EK. PPtree: Projection pursuit classification tree. Electronic Journal of Statistics. 2013;7(1):1369–1386. doi:10.1214/13-EJS810.

40. Ein-Dor L, Zuk O, Domany E. Thousands of samples are needed to generate a robust gene list for predicting outcome in cancer. Proceedings of the National Academy of Sciences of the United States of America. 2006;103(15):5923–5928. doi:10.1073/pnas.0601231103.

41. Chen C, Liaw A, Breiman L. Using Random Forest to Learn Imbalanced Data. University of California, Berkeley. 2004; p. 1–12.

42. Saito T, Rehmsmeier M. The Precision-Recall Plot Is More Informative than the ROC Plot When Evaluating Binary Classifiers on Imbalanced Datasets. PLoS ONE. 2015; p. 1–21. doi:10.1371/journal.pone.0118432.

43. Weng CG, Poon J. A New Evaluation Measure for Imbalanced Datasets. Proceedings of the 7th Australasian Data Mining Conference. 2008;87.

44. Floares AG, Ferisgan M, Onita D, Ciuparu A, Calin GA, Manolache FB. The Smallest Sample Size for the Desired Diagnosis Accuracy. International Journal of Oncology and Cancer Therapy. 2017;2.

45. He Z, Yu W. Stable feature selection for biomarker discovery. Computational Biology and Chemistry. 2010;34(4):215–225. doi:10.1016/j.compbiolchem.2010.07.002.

46. Han Y, Lei Y. A variance reduction framework for stable feature selection. Statistical Analysis and Data Mining: The ASA Data Science Journal. 2012;8(5):428–445. doi:https://onlinelibrary.wiley.com/doi/epdf/10.1002/sam.11152.

47. Janitza S, Hornung R. On the overestimation of random forest’s out-of-bag error. PLoS ONE. 2018;13(8):1–31. doi:10.1371/journal.pone.0201904.

48. Toloşi L, Lengauer T. Classification with correlated features: Unreliability of feature ranking and solutions. Bioinformatics. 2011;27(14):1986–1994. doi:10.1093/bioinformatics/btr300.

49. Statnikov A, Aliferis CF. Analysis and computational dissection of molecular signature multiplicity. PLoS computational biology. 2010;6(5):e1000790.

50. Wang H, Yang F, Luo Z. An experimental study of the intrinsic stability of random forest variable importance measures. BMC Bioinformatics. 2016; p. 1–18. doi:10.1186/s12859-016-0900-5.

51. Deng H. Interpreting tree ensembles with inTrees. International Journal of Data Science and Analytics. 2019;7(4):277–287. doi:10.1007/s41060-018-0144-8.

52. Risso D, Schwartz K, Sherlock G, Dudoit S. GC-Content Normalization for RNA-Seq Data. BMC Bioinformatics. 2011;12(1):480.

53. Saeys Y, Abeel T, Yves VdP. Feature selection techniques for maximum entropy based biomedical named entity recognition. In: ECML PKDD 2008, Part II, LNAI 5212; 2008. p. 313–325.

