## Supplementary figures and images for "Assessing Random Forest self-reproducibility for optimal short biomarker signature discovery"

### S1 Fig

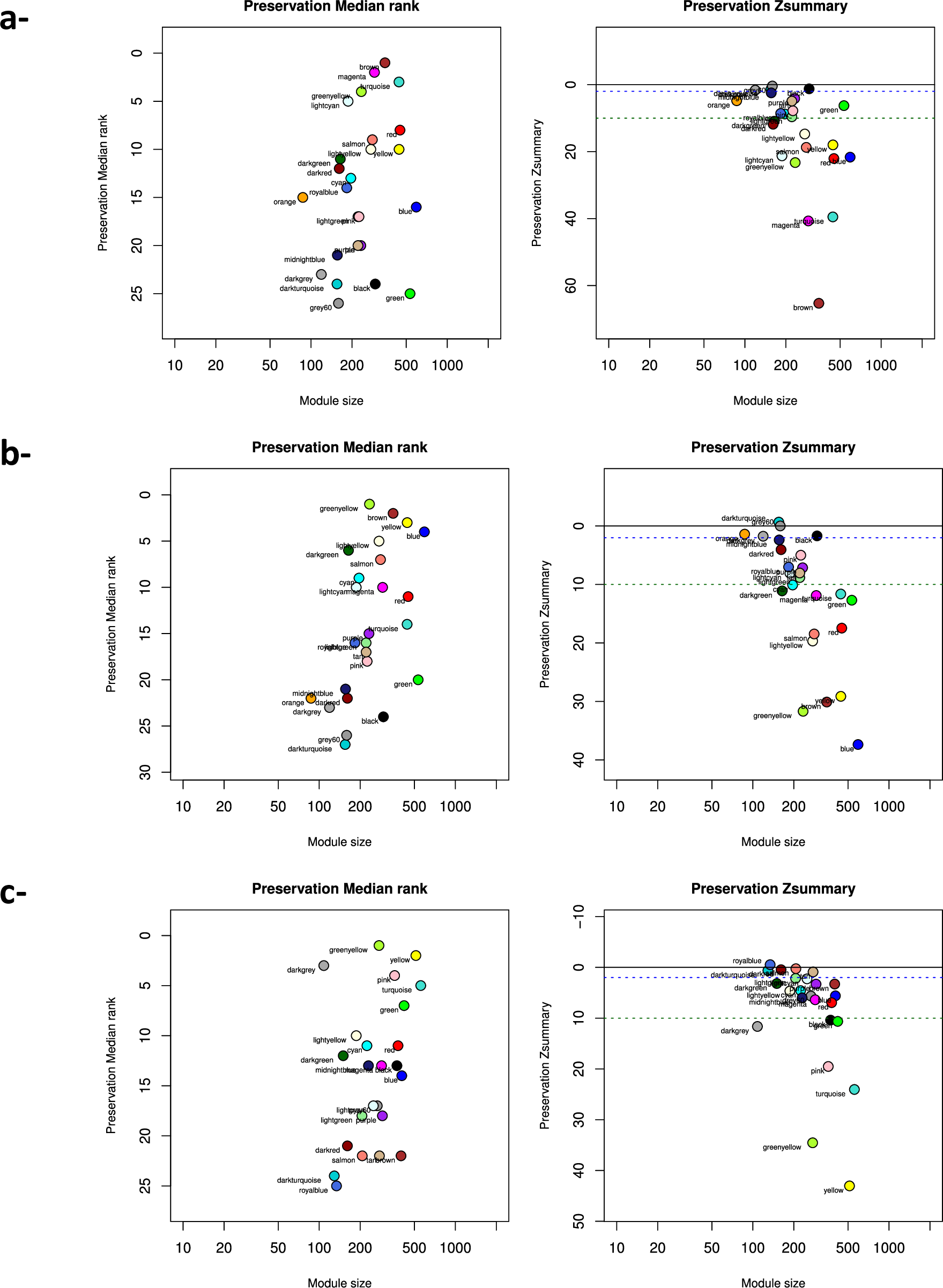

### S2 Fig

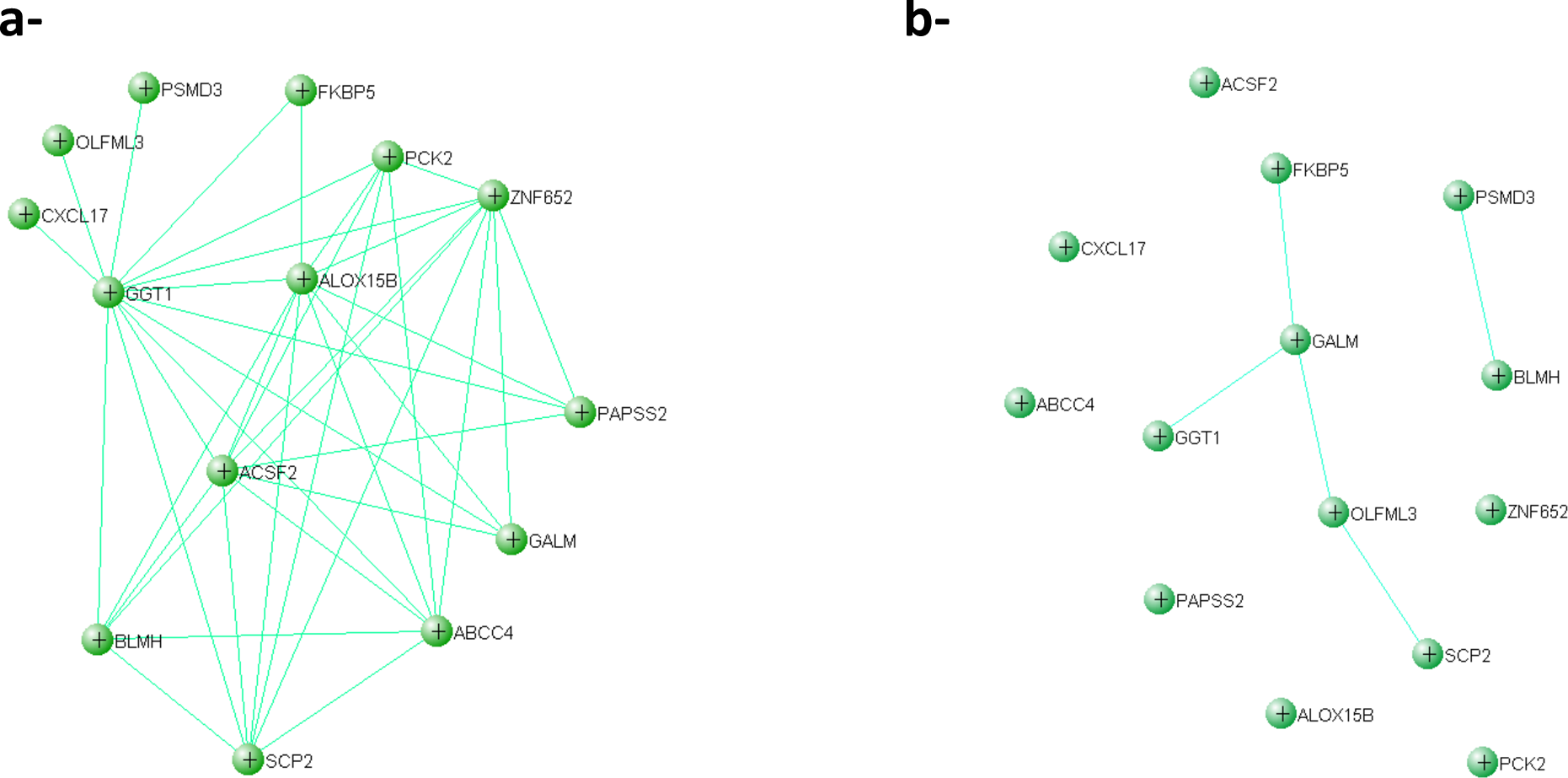

### S3 Fig

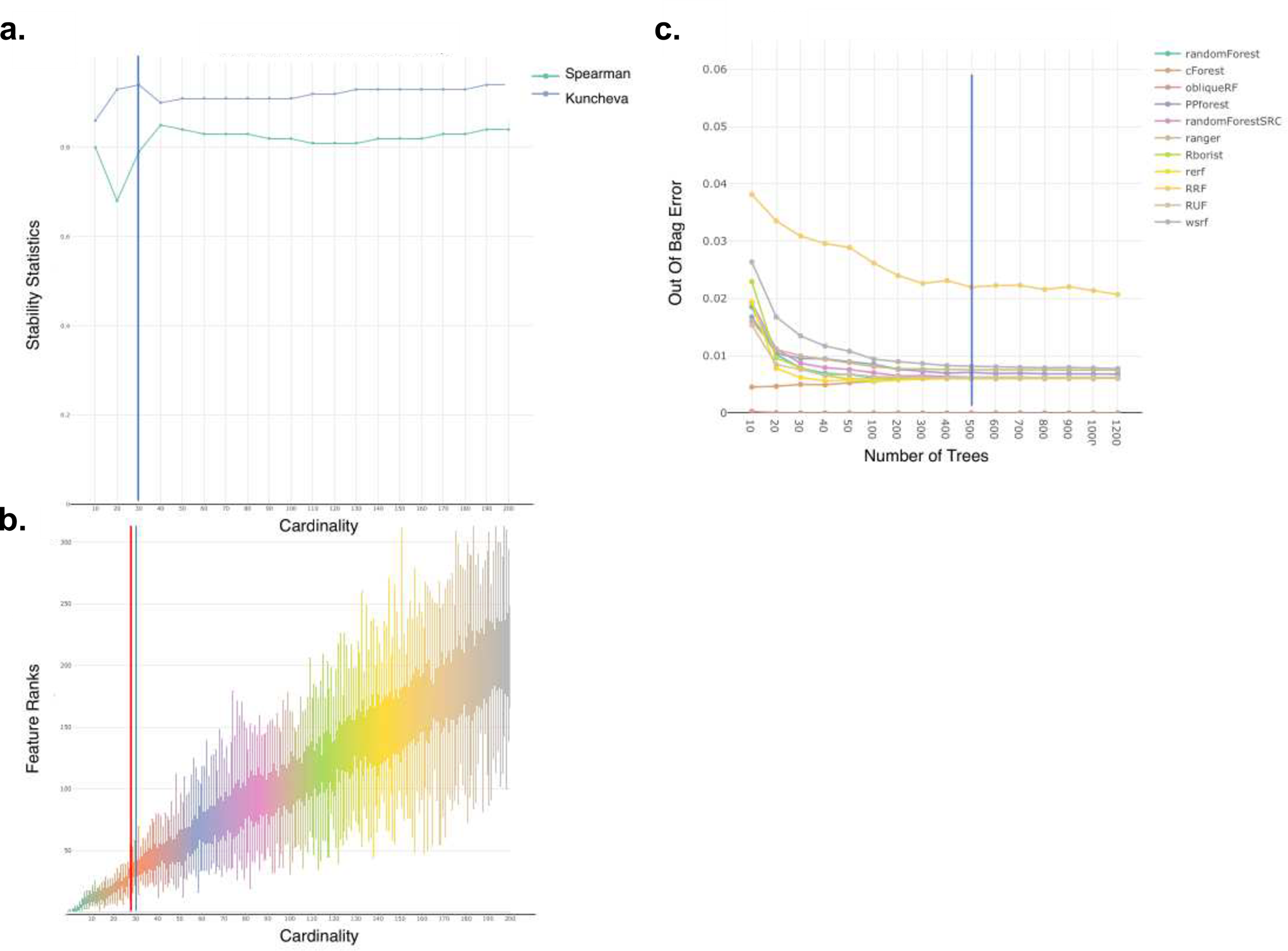

### S4 Fig

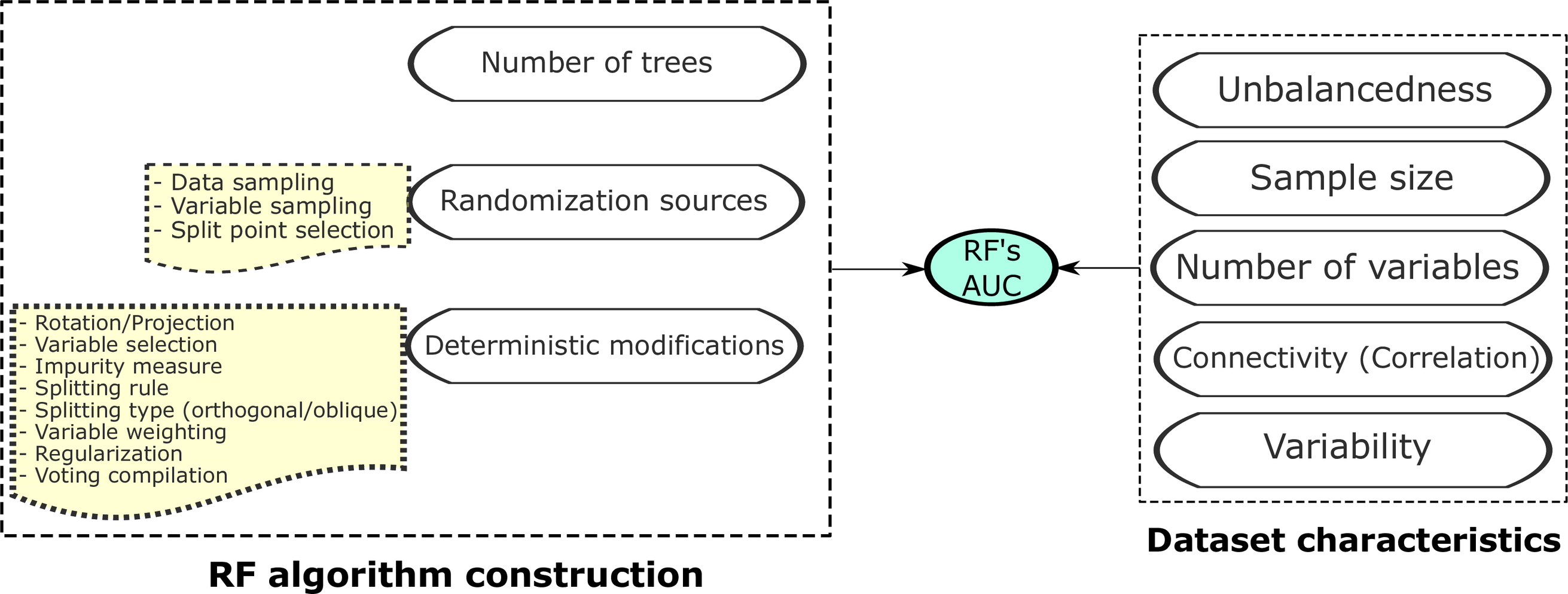

### S5 Fig

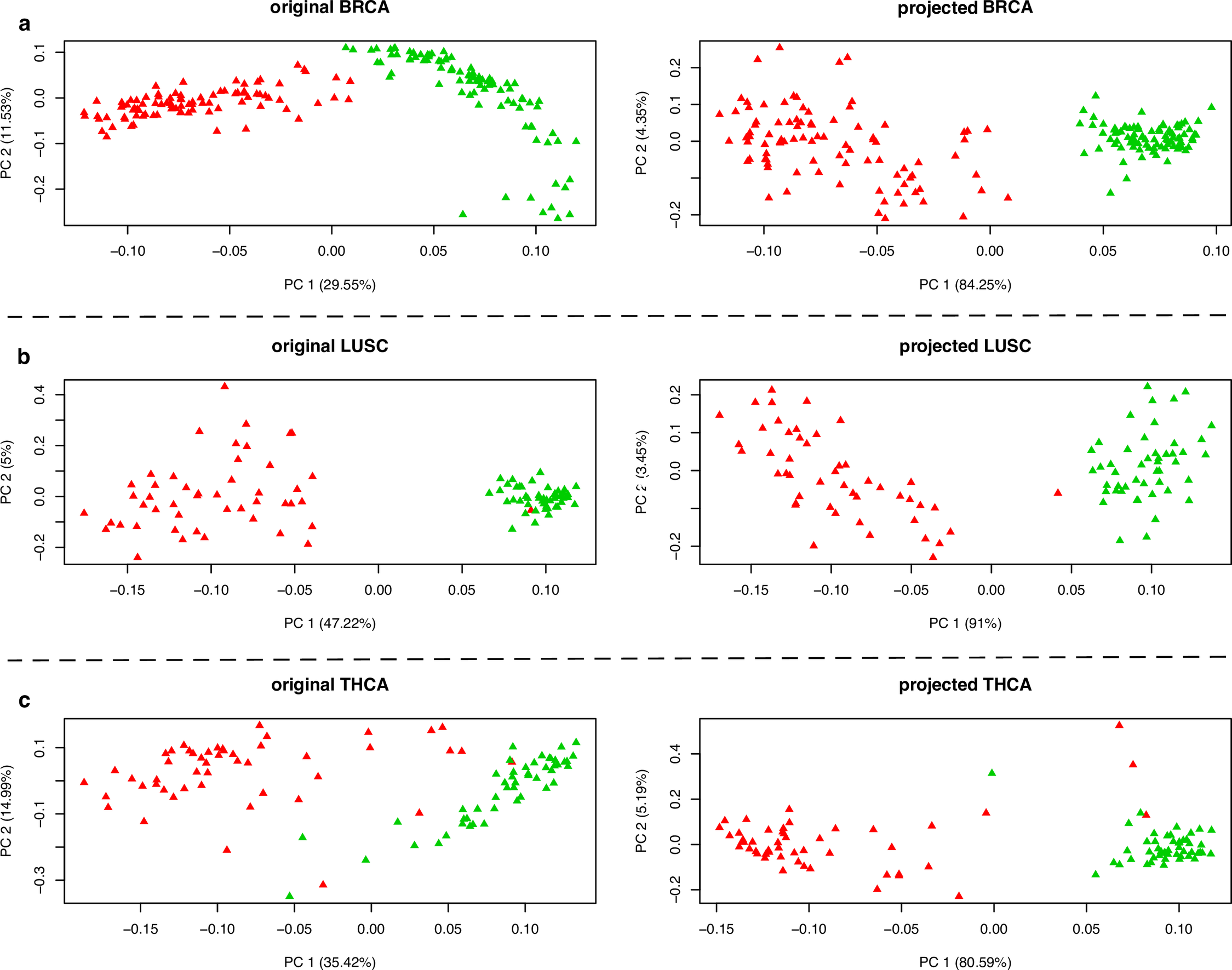
