## Supplementary material for "Assessing Random Forest self-reproducibility for optimal short biomarker signature discovery": S1 File

### Supplementary Methods

#### Stable RF-based Feature Selection

A total of $k = 50$ balanced partitions were randomly defined from the original TCGA datasets, using a sampling perturbation rate $p = 0.9$. For each random partition, $q = 25$ RF model importance were calculated using the randomForest R-package with the default parameters [11]. The Mean Decrease in Accuracy $MDA$ and the Mean Decrease in Gini $MDG$ were computed for all the variables over the $q$ models. The $MDA$ and $MDG$ were respectively ranked, and their average rank was used to rank the variables for each partition. The $k$ ranking sequences obtained were used to assess the stability of the first 200 variables using the Spearman (correlation) and the Kuncheva (overlap) statistics from the R-package OmicsMarkeR The first local maxima, common between Spearman and Kuncheva, gave the minimal set of important variables $Nv'$. Good stability values (i) were obtained for the Kuncheva index ($i \geq 0.8$), which means the models selected almost the same variables at each run [50].

#### Assessing RF randomness within identical AUCs

To assess whether the RF implementations keep their intrinsic randomness, we extracted the rules of two distinct models that produced the same AUC. The R package "inTrees" was used to extract these rules [51]. Table S3 Table displays an example of the rules from two randomly selected randomForest models, trained on one signature-resampling combination. A more detailed example on each dataset is also available in S4 Table, which displays the rules from two random randomForest models, trained on three signature-resampling combinations. The genes, the thresholds, and the number of steps used were different in the three rules and the two models, while the AUC was precisely 1. These results indicated that the inherent randomness was conserved in our methodology.

### Supplementary results

#### Short signature design

**Feature Selection and number of trees.** To keep the minimal amount of important variables at $Nv' < 50$ for the purpose of the short BSD, we computed the Kuncheva and Spearman indexes from the variance-filtered TCGA datasets. On the top 200 most important variables, the cardinality corresponding to the first local maxima of Kuncheva and Spearman indexes were matched to the rank distribution of each variable (S3 Fig a and b, and $Nv'$ in S1 Table). Finally, we assessed the number of trees to use in the modelizations. We set it as the common number of trees that reached a minimal and stable OOB error between RF methods, and $Nt = 500$ was obtained (S3 Fig c). The parameters selected for this step were summarized in S1 Table.

**Efficiency of the feature selection used.** To assess whether the features selected with the FS led to the separation of the tumor and healthy classes, we applied a PCA on each dataset. Using the plotPCA function of the EDASeq package [52], For BRCA (S5 Fig a), 89% of the variability was explained by the two first components after the FS, while only 50% was explained on the original dataset. For all three datasets, the FS increased the variability captured by the two first components when compared to the original datasets. Nevertheless, four samples of the THCA dataset were not well clustered after the FS. Consequently, except for a few resistant THCA samples, the features selected could separate the samples according to their expected category.

**Correlation between the variables within a signature.** To understand how the combinations of the selected features may bias the modelizations, we calculated the correlation between each signature's variables. Therefore, we looked at the overall correlations observed between the variables and the percentage of highly-correlated variables within each signature (see S1 Table). We calculated the overall correlation as the average Pearson correlation between the variables selected by the FS in the whole dataset. We set the percentage of highly-correlated variables within a signature as the proportion of signatures for which the variables achieved the Pearson correlation $\geq 0.75$ over all the resamplings. The LUSC dataset displayed strong variable correlations within each signature and for most of the resamplings. On the opposite, only a few strong correlations were observed for the BRCA and THCA datasets. Interestingly, these strong correlations stuck to few resamplings. By resetting the signature set with new signature combinations, similar patterns of correlation were observed.

#### Supplementary Discussion

The FS developed in the current methodology was mostly based on a previous observation made by Alelyani *et al.* [27]. This FS aimed at capturing the most stable variables among thousands from each dataset.

Gene expression datasets with small-size samples and high dimensional features were previously described as suffering from intrinsic instability [27, 45, 50]. Interestingly, we did not observe such an effect on the three datasets, maybe the prior filtering based on paired tumor / healthy samples.

The Kuncheva (overlap) and Spearman (correlation) indexes were used since we expected high ranking correlations with such a number of variables. The Spearman index was chosen over the Pearson index because of its compatibility with feature weighting [45, 53]. The minimal number of stable features was set at the first local maxima reached by both indexes as proposed by Alelyani *et al.* [27] and refined by discarding variables with a large dispersion of their ranking (S3 Fig b). Such visual selection could influence the number of variables obtained, and subsequently, this step should be improved for a more objective choice.

The FS used the variable importance measures MDG and MDA of the original RF algorithm described by Breiman in 2001 [10] as a standard. Whereas the randomForest package was used for the FS, our results did not rank it as the best algorithm. With 10 of the 15 RF methods implementing a FS algorithm we expected up to ten different selected features to start our comparisons, which would have been hard to handle. Nevertheless, this study would benefit from results obtained based on the nine other FS. Besides, the FS used a data perturbation of 0.9 together with an average of 50 rounds of FS. While other perturbations were possible, we aimed at reaching a set with the most stable variables for each dataset with an average AUC close to 1. Such a high average AUC was mandatory to demonstrate the differences in AUC hyper-stabilities for the purpose of the current study.
