## Supplementary material for "Assessing Random Forest self-reproducibility for optimal short biomarker signature discovery": S1 Table

| Symbol | Description |
|---|---|
| $Ns$ | Number of all possible combinations (signatures) generated from $Nv'$ variables |
| $Nv$ | Number of variables of TCGA datasets after the normalization and filtering step |
| $Nv'$ | Minimum number of variables to be kept for downstream analysis. This number is determined by FS and stability indices. |
| $Nt$ | Finetuned value of *ntree* parameter for all RF implementations. |
| $S$ | Number of different sized signatures randomly selected from the set of $Ns$ signatures |
| $p$ | Resampling rate. It refers to the percentage of data that goes to training partition after a balanced random sampling from the original data. |
| $k$ | Number of partitions randomly sampled from the dataset |
| $q$ | Number of intrinsic RF models constructed/used for each partition-signature combination, and for each RF implementation |
| $RFr$ | Total number of runs of each RF implementation |
| $CV$ | Coefficient of variation of the $q = 25$ AUC values generated for each partition-signature combination, and for each RF implementation |
| $s$ | Sample standard deviation |
| $\bar{x}$ | and $\bar{x}$ Sample mean |
| $S_0$ | Number of signatures with $CV == 0$ for a given resampling partition |
| $k_0$ | Number of resampling partitions with $CV == 0$ for a given signature |
| $HR_{k_n}$ | Hyper-stability of the resampling $k_n$ |
| $HS_{S_n}$ | Hyper-stability of the signature $S_n$ |
| $HRS$ | Mean of non-zero $HRs$ across all resampling partitions |
| $HSS$ | Mean of non-zero $HSs$ across all signatures |
