## Supplementary material for "Assessing Random Forest self-reproducibility for optimal short biomarker signature discovery": S2 Table

| Dataset | TCGA-BRCA | TCGA-LUSC | TCGA-THCA |
|---|---|---|---|
| Histological SubType | infiltrating ductal carcinoma | lung squamous cell carcinoma | thyroid papillary carcinoma |
| Patients | 91 | 48 | 49 |
| Samples | 182 | 96 | 98 |
| $Nv$ | 9560 | 9262 | 9353 |
| $Nv'$ (FS Stability) | 28 | 9 | 38 |
| $Nt$ (OOBerr) | 500 | 500 | 500 |
| Selected Signatures | 78 | 21 | 108 |
| Overall Variable Correlation | 0.61 | 0.83 | 0.64 |
| Highly-correlated variables within-signature (%) | 0.02 | 0.96 | 0.03 |
| Variable-Sample Ratio | 0.3 | 0.2 | 0.8 |
