## Supplementary material for "Assessing Random Forest self-reproducibility for optimal short biomarker signature discovery": S4 Table

| rule/step | signature | resampling | model | condition | prediction |
|---|---|---|---|---|---|
| 1/1 | s3 | r47 | model1 | PTPRB>2385.9889 | normal |
| 1/2 | s3 | r47 | model1 | Else | tumor |
| 1/1 | s3 | r47 | model2 | GPIHBP1<=451.4196 | tumor |
| 1/2 | s3 | r47 | model2 | Else | normal |
