## Supplementary material for "Assessing Random Forest self-reproducibility for optimal short biomarker signature discovery": S5 Table

| dataset | step | rule | signature | resampling | model | length | frequency | error | condition | prediction |
|---|---|---|---|---|---|---|---|---|---|---|
| BRCA | 1 | 1 | signature2 | resampling1 | model1 | 2 | 0.50 | 0 | COL10A1<=899.8924 & SDPR>543.7528 | normal |
| BRCA | 2 | 1 | signature2 | resampling1 | model1 | 1 | 0.50 | 0 | Else | tumor |
| BRCA | 1 | 1 | signature2 | resampling1 | model2 | 2 | 0.50 | 0 | COL10A1>84.6616 & HLF<=648.1244 | tumor |
| BRCA | 2 | 1 | signature2 | resampling1 | model2 | 1 | 0.50 | 0 | Else | normal |
| BRCA | 1 | 2 | signature30 | resampling48 | model1 | 2 | 0.50 | 0 | EZH1>983.8384 & GPRASP1>278.3492 | normal |
| BRCA | 2 | 2 | signature30 | resampling48 | model1 | 1 | 0.50 | 0 | Else | tumor |
| BRCA | 1 | 2 | signature30 | resampling48 | model2 | 1 | 0.48 | 0 | CALCOCO1>2232.891 | normal |
| BRCA | 2 | 2 | signature30 | resampling48 | model2 | 1 | 0.51 | 0.02 | EZH1<=1269.0316 | tumor |
| BRCA | 3 | 2 | signature30 | resampling48 | model2 | 1 | 0.01 | 0 | Else | normal |
| BRCA | 1 | 3 | signature70 | resampling21 | model1 | 1 | 0.50 | 0 | MMP11<=406.0273 | normal |
| BRCA | 2 | 3 | signature70 | resampling21 | model1 | 1 | 0.50 | 0 | Else | tumor |
| BRCA | 1 | 3 | signature70 | resampling21 | model2 | 1 | 0.50 | 0 | PPP1R12B<=2025.1838 | tumor |
| BRCA | 2 | 3 | signature70 | resampling21 | model2 | 1 | 0.50 | 0 | Else | normal |
| LUSC | 1 | 1 | signature3 | resampling47 | model1 | 1 | 0.50 | 0 | PTPRB>2385.9889 | normal |
| LUSC | 2 | 1 | signature3 | resampling47 | model1 | 1 | 0.50 | 0 | Else | tumor |
| LUSC | 1 | 1 | signature3 | resampling47 | model2 | 1 | 0.50 | 0 | GPIHBP1<=451.4196 | tumor |
| LUSC | 2 | 1 | signature3 | resampling47 | model2 | 1 | 0.50 | 0 | Else | normal |
| LUSC | 1 | 2 | signature11 | resampling27 | model1 | 1 | 0.50 | 0 | TGFBR2>7580.5877 | normal |
| LUSC | 2 | 2 | signature11 | resampling27 | model1 | 1 | 0.50 | 0 | Else | tumor |
| LUSC | 1 | 2 | signature11 | resampling27 | model2 | 1 | 0.50 | 0 | GPIHBP1>451.4196 | normal |
| LUSC | 2 | 2 | signature11 | resampling27 | model2 | 1 | 0.50 | 0 | Else | tumor |
| LUSC | 1 | 3 | signature21 | resampling3 | model1 | 1 | 0.50 | 0 | GPR116<=7183.4553 | tumor |
| LUSC | 2 | 3 | signature21 | resampling3 | model1 | 1 | 0.50 | 0 | Else | normal |
| LUSC | 1 | 3 | signature21 | resampling3 | model2 | 1 | 0.50 | 0 | ESAM>2673.1366 | normal |
| LUSC | 2 | 3 | signature21 | resampling3 | model2 | 1 | 0.50 | 0 | Else | tumor |
| THCA | 1 | 1 | signature14 | resampling5 | model1 | 2 | 0.50 | 0 | DLG4>381.1242 & EPHB1<=570.4257 | tumor |
| THCA | 2 | 1 | signature14 | resampling5 | model1 | 1 | 0.50 | 0 | Else | normal |
| THCA | 1 | 1 | signature14 | resampling5 | model2 | 2 | 0.50 | 0 | AGPAT4>93.7298 & C6orf168<=99.0904 | normal |
| THCA | 2 | 1 | signature14 | resampling5 | model2 | 1 | 0.50 | 0 | Else | tumor |
| THCA | 1 | 2 | signature65 | resampling23 | model1 | 2 | 0.50 | 0 | GALE<=498.265 & ODZ1>19.2609 | normal |
| THCA | 2 | 2 | signature65 | resampling23 | model1 | 1 | 0.50 | 0 | Else | tumor |
| THCA | 1 | 2 | signature65 | resampling23 | model2 | 2 | 0.50 | 0 | METTL7B<=291.4223 & ODZ1>19.2609 | normal |
| THCA | 2 | 2 | signature65 | resampling23 | model2 | 1 | 0.50 | 0 | Else | tumor |
| THCA | 1 | 3 | signature101 | resampling42 | model1 | 2 | 0.50 | 0 | FHOD1<=876.5515 & SRCIN1<=210.8955 | normal |
| THCA | 2 | 3 | signature101 | resampling42 | model1 | 1 | 0.50 | 0 | Else | tumor |
| THCA | 1 | 3 | signature101 | resampling42 | model2 | 2 | 0.50 | 0 | MCTP2>340.0583 & METTL7B<=289.4312 | normal |
| THCA | 2 | 3 | signature101 | resampling42 | model2 | 1 | 0.50 | 0 | Else | tumor |
