## Supplementary material for "Assessing Random Forest self-reproducibility for optimal short biomarker signature discovery": S6 Table

| Method | Min | Max | % Stable |
|---|---|---|---|
| ccf | 0 | 1.11 | 61.24 |
| cForest | 0 | 1.15 | 70.95 |
| extraTrees | 0 | 1.01 | 74.67 |
| iForest | 0 | 1.06 | 67.24 |
| obliqueRF | 0 | 1.07 | 91.81 |
| PPforest | 0 | 31.51 | 31.33 |
| randomForest | 0 | 1.04 | 68.29 |
| randomForestSRC | 0 | 0.09 | 93.90 |
| randomUniformForest | 0 | 0.17 | 74.10 |
| ranger | 0 | 1.08 | 67.90 |
| Rborist | 0 | 0.57 | 77.81 |
| rerf | 0 | 1.15 | 62.10 |
| rotationForest | 0 | 1.78 | 55.43 |
| RRF | 0 | 2.03 | 11.62 |
| wsrf | 0 | 0.09 | 94.29 |
